# The minimum intervention principle of optimal control relates the uncontrolled manifold to muscle synergies

**DOI:** 10.1101/2023.08.18.553939

**Authors:** Neelima Sharma

## Abstract

The nervous system uses muscle activation patterns to perform successful motor tasks, and motor tasks inevitably satisfy the laws of mechanics. The number of muscles usually exceeds the degrees of freedom of the task such that multiple combinations of muscle activities satisfy mechanical demands. A low-dimensional space usually explains a large variance in the muscle activations, leading to the hypothesis that muscle redundancy is solved by neurally coordinated muscle synergies. In addition to synergies, motor task mechanics also enforce a structure on muscle activity. Muscles satisfy multiple non-negotiable mechanical demands of equilibrium, stability, force, and respect the constraints. The redundancy in the muscle architecture is the degree of freedom available after accounting for the necessary mechanics. In this study, I investigate how task mechanics structure muscle activities by using a biomechanical model of an index finger in contact and published measurements of seven muscle activities during a fingertip force production task. I derive a map from muscle activities to complete task mechanics with the necessary conditions of equilibrium, stability, and force. By invoking the uncontrolled manifold hypothesis, I show that the variability in muscle activities is channeled in the task-irrelevant directions, which is given by the null space of the map from muscle activations to task variables. Furthermore, I show that the principal component that explains maximum variance in muscle activations is oriented along the task-irrelevant direction with the highest projected variance, suggesting that the maximal principal directions correspond to the task-irrelevant subspace rather than the task-relevant directions. This study has consequences for understanding muscular redundancy and synergy, and also provides direct evidence that the simplified biomechanical models satisfy mechanical requirements.

## I. Introduction

Flipping a page or scrolling a screen requires precisely controlled fingertip forces in addition to a stable finger posture that maintains equilibrium. So muscle activities satisfy multiple mechanical demands to accomplish successful motor tasks. The number of muscles generally exceeds the number of mechanical degrees of freedom of the task variables, such as the finger posture and tip forces, leading to infinite possibilities for selecting muscle activities. How such muscular redundancy [1] is resolved is debated. Examples of mechanisms invoked to address muscle redundancy are muscle synergies [2–5], robustness to muscle dysfunction [6], optimization of physiological effort [7, 8], and control of motor task mechanics [9, 10]. Muscle synergies are a hypothesized level of control exerted by the central nervous system where group of muscles are controlled in synergy rather than as independent muscle units, thus reducing the dimensions of the control variable. But mechanics of motor task also impose a structure on the muscle activations with consequences for interpreting the degree of redundancy. In this study, I focus on how task mechanics impose a structure on muscle activations, and how the structure compares with the proposed definitions of muscle synergies.

A way to obtain the structure imposed by mechanics on muscle activations is by applying the uncontrolled manifold (UCM) hypothesis [11]. In the context of a muscle-driven motor behavior, UCM hypothesis states that the noise associated with muscle activities is channeled in the task-irrelevant subspace and reduced in the task-relevant subspace of the control variables (e.g. muscle activities). By channeling the noise in the task irrelevant directions of the high-dimensional control variable, the task variables (e.g. limb position or tip forces) are minimally affected [11–14]. By yielding the control of variables within the uncontrolled manifold (task-irrelevant directions), motor behavior follows the minimal intervention principle of optimal control [15, 16]. For example, during the task of producing instructed fingertip forces with the index finger, the noise in muscle activities is channeled in the uncontrolled manifold of the map from muscle activations to fingertip forces [14], but other studies show that task-irrelevant space is also controlled [17]. To calculate the true degree of redundancy for the static task of producing fingertip forces, one must include equilibrium and stability of the finger posture in the description of the task as they are necessary mechanical demands that impose additional structure on muscle activities.

In this study, I test how the complete description of motor task mechanics that includes the non-negotiable demands of equilibrium, force, stability, and constraint satisfaction affect muscular redundancy. I use published data on the activities of the seven muscles of an index finger during the fingertip force production task [14] and a mathematical model of the index finger [18] to analyze the structure of variability in muscle activities. I test whether the uncontrolled manifold hypothesis applies by calculating the variability channeled in the uncontrolled subspace of the map from muscle activations to the necessary mechanical needs. Previous studies use principal component analysis of muscle activities to show that a low dimensional activation subspace typically explains most of the variance and interpret the maximal principal directions as evidence of muscle synergies that depict the few task-relevant directions [19, 20]. In contrast, the uncontrolled manifold hypothesis states that variance is channeled in the task-irrelevant space of the mapping from the control to the task variable [11]. So, I hypothesize that the high variance subspace calculated from the principal component analysis must reflect the uncontrolled task-irrelevant null space, and calculate the alignment of the null space with the maximal principal direction. Whether muscle synergies exist and are aware of the structure imposed by mechanics, and if synergy sets are so organized as to facilitate the motor task at hand, remains a goal of future investigations.

## II. Methods

### A. Data

I analyze the data used in the study ‘Structured Variability of Muscle Activations Supports the Minimal Intervention Principle of Motor Control’ by Valero-Cuevas et al., 2009 [14]. The study recorded the muscle activations from all the seven muscles that drive the index finger while the index finger produces instructed fingertip forces. In the experiment, volunteers held a dowel attached to the ground and produced instructed fingertip forces on the surface of a load cell using their index fingers. The posture of the finger had neutral abduction and proximo-distal joint angles of 30°, 45°, and 15° flexion at the metacarpophalangeal, proximal interphalangeal, and distal interphalangeal joints, respectively. The instructed force pattern consisted of an initial hold phase of 2 N for 4 s, followed by a randomly varying phase of 10 s, which was followed by a final hold phase of 10 s. During each phase, muscle electromyograms were recorded from all seven muscles that drive the index finger using fine-wire electrodes. I use data from 6 subjects with a total of 35 trials. For additional details on data collection, see [14].

#### Signal processing

EMG recordings were band-stop filtered in the range 58 Hz to 62 Hz and 118 Hz to 122 Hz for set C to remove the electrical artifacts, followed by high-pass filtering at 20 Hz to remove movement artifacts, full-wave rectified, and passed through fourth-order Butterworth filters with time constants of 0.03 s and 0.23 s to adjust for the muscle’s excitation-contraction dynamics [14].

### B. Index Finger Model

To find the relationship between muscle activations and the conditions of force, equilibrium, and stability, I analyze a mathematical model of the index finger driven by seven muscles. The model, its dynamics and stability analysis are completely derived in a previous study [18]. In this study, the vector of configuration dependent forces and torques are ignored (see II E for the rationale). I provide a brief summary of the pertinent model features for the current study.

#### Dynamics of the index finger

An index finger in contact is modeled as a planar three link manipulator, with contact constraints at the endpoints. The dynamics of the index finger are given in terms of the mass matrix of the linkage network M *∈* ℝ^3×3^, the Coriolis and centripetal contribution C *∈* ℝ^3×3^, the vector of joint torques and forces 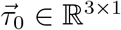, the stiffness matrix K *∈* ℝ^3×3^, the damping matrix B *∈* ℝ^3×3^, and the vector of neutral joint angles 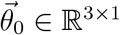 (Eq. (1), [18]). The kinematic constraint of endpoint contact is described as a constant position vector 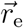, rewritten in terms of a posture dependent Jacobian J *∈* ℝ^*p×n*^, and the vector of Lagrange multiplier 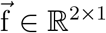 that expresses the forces due to the constraint (Eq. (1), [18]). The dynamical equations, constraint equation, and the constraint forces, respectively, are:

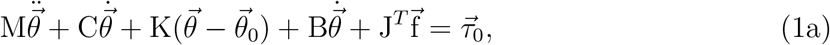

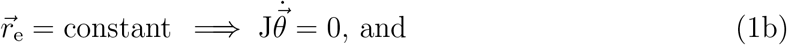

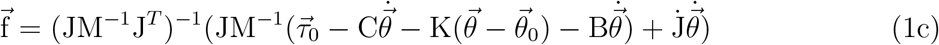

Equilibrium at the posture 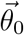 requires that the joint torques 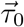 support the endpoint force 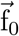. By setting 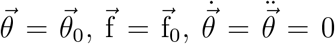 in equation (1a), and using 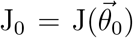, the joint torques are obtained as:

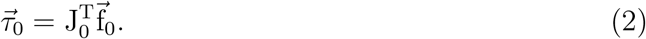

By linearizing the governing equations and solving for marginal stability of the system, the minimum stiffness matrix for stability is given as (see [18] for complete derivation):

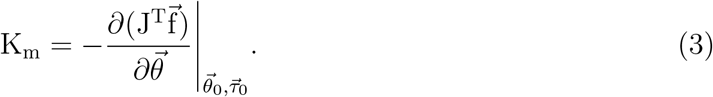

#### Muscle model

The activations of the seven muscles of the index finger (table S1) are parameterized by the activation vector 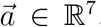. Muscle activites map to muscle tension that lead to joint torques. The mapping depends on the diagonal matrix F_iso_ *∈* ℝ^7×7^ of maximal isometric force of each muscle, and the matrix R *∈* ℝ^3×7^ of moment arms of each musculotendon about each joint. In terms of these parameters, the vector of joint torques 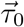 is given by,

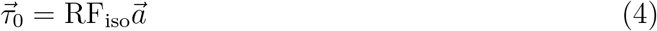

Muscles contraction also gives rise to stiffness at the index finger joints. Whether the finger posture remains passively stable or not depends on whether the muscle induced stiffness is greater than the minimum stabilizing stiffness required for stability [18]. Our previous study [18] shows that short-range muscle stiffness ensures the stability of the finger posture against buckling while producing maximal fingertip forces. Here, I test whether humans also rely on short-range muscle stiffness for sub-maximal forces. However, I do not compare the muscle induced stiffness with the minimal condition in this study but test a different hypothesis. For a one degree of freedom system such as a planar finger in contact, the minimum stabilizing stiffness matrix has only one non-zero eigenvalue with the corresponding eigenvector given by a specific combination of joint angles, that are determined by the fingertip constraint. I test whether muscle activities predominantly stiffen the finger in the direction of the pertinent eigenvector, which reflects the degree of freedom of the system, in comparison to the eigendirections that are automatically fixed due to the tip constraint and thereby do not require additional stiffening for stability.

### C. Muscle activations satisfy task mechanics

The success of the motor task of producing desired fingertip forces relies on selecting muscle activations that simultaneously satisfy the mechanical requirements of force, equilibrium, and stability. I derive the transformation maps associated with each condition below and combine them to derive the complete description of the task-relevant space.

#### Condition of desired force

Valero-Cuevas et al. (2009) [14] showed that the variability in muscle activations is channeled in the task-irrelevant subspace of the mapping from muscle activations to fingertip forces during a static task. Although the subjects are instructed to produce finite distal forces, lateral and palmar fingertip forces need to respect the friction cone to avoid slipping, and therefore, all three components of the force vector are controlled [14]. Here, I repeat the analysis. A linear model is used to map the muscle activations to the tip force vector:

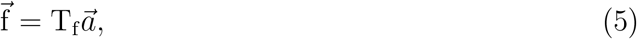

where T_f_ *∈* ℝ^3×7^ is a time-invariant transformation matrix that maps normalized muscle tensions to fingertip forces [21]. For T_f_, there exists a 4D task-irrelevant null-space and 3D task-relevant complement to the null-space. Complete trials are used for calculating the force map. The R^2^ for distal fingertip force for the computed T_f_ is 0.88 ± 0.07 (mean ± standard deviation). Monte Carlo simulations are used to verify that linear fitting does not automatically channel the variance in the null space (S2).

#### Condition of static equilibrium

Using the equilibrium condition Eq. (2) and the relationship Eq. (1c) between joint torques and the fingertip force, the condition for an activation vector 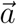 to maintain equilibrium, i.e. 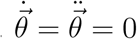, is given by

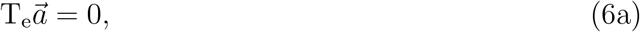

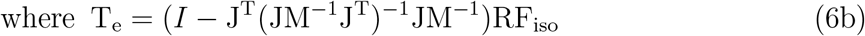

Thus, the muscle activation vector satisfies equilibrium only when it belongs to the null-space of T_e_ *∈* ℝ^3×7^. For the index finger model, the matrix (I *−* J^T^(JM^*−*1^J^T^)^*−*1^JM^*−*1^) is a projection matrix with two zero eigenvalues or one eigenvalue equal to one. So there exists a six-dimensional null-space of T_e_. Variability in the null-space of T_e_ does not affect equilibrium, but any variability in the codimension-1 row-space of T_e_ leads to a loss of equilibrium. So, there exists a 6D task-irrelevant subspace and 1D task-relevant subspace.

#### Condition for stability

The task relevant manifold for stability is calculated by finding the map from the muscle activation vector to the minimum stabilizing stiffness matrix. Using equations (1c), (3) and imposing equilibrium, i.e. 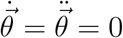, I obtain

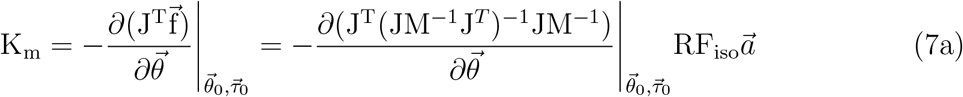

The stiffness matrix K_*m*_ is characterized by one finite eigenvalue ζ along the null space to the tip constraints, 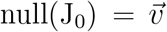. I project the stiffness matrix on the null space to the constraints by using the projection matrix 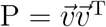 to obtain a map from 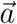 to the relevant stiffness eigendirection (S1):

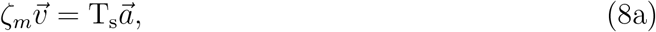

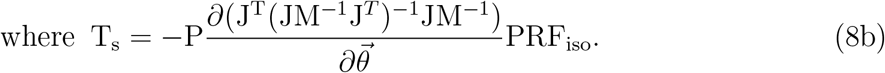

Here, T_s_ *∈* ℝ^3×7^ is time-invariant matrix that has a 6D task-irrelevant null space and a 1D task-relevant complement space. The null space of T_s_ defines the subspace where variations in muscle excitations do not change the postural stiffness in the pertinent eigendirection.

#### Combined equilibrium and force condition

The map from muscle activations to the task variables for the task consisting of equilibrium and force conditions is given by T_*ef*_. When both equilibrium and force requirements are considered in the task description, the task-irrelevant subspace is given by:

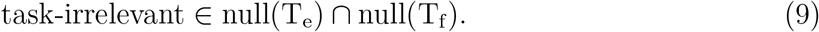

#### Complete task description

A complete task description includes equilibrium, force, and stability requirements, with the map from muscle activations to task mechanics given by T_efs_. The intersection of force, equilibrium, and stiffness null spaces gives the complete task-irrelevant subspace:

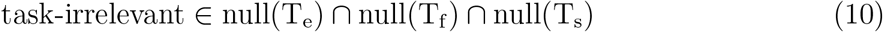

### D. Analysis of variability patterns

#### a. Calculation of task-relevant and irrelevant variability

I calculate the full covariance matrix of within and cross muscle interaction G_*n*_ during the final constant phase of each trial, as done previously [14]. The final hold phase is used because it is the longest and subjects are expected to have settled on their motor control strategy. Next, I project the covariance matrix onto the task-irrelevant and relevant subspaces, provided by the null-space and the complement to the null-space of the biomechanical transformation matrix, respectively. The transformation matrices considered were T_e_ for equilibrium, T_f_ for force, T_s_ for stability, T_ef_ for combined equilibrium and force condition, and T_efs_ for the complete task description containing equilibrium, force, and stability condition.

To test the hypothesis that equilibrium, stiffness, and force constraints are associated with variance attenuation in the task-relevant subspace, an orthonormal basis for the null-space and the complement space for each of the transformation matrices is used, the variance is projected on each orthonormal axis, and a task-relevant and task-irrelevant average is calculated by using the number of null and complement basis vectors [14]. I computed the variability ratio *η* by dividing the task-relevant variability by the task-irrelevant variability. When this ratio is significantly lower than 1, there exists support for the hypothesis that muscle activations support the minimum intervention principle. For computing the null-space of the complete task transformation map, I used a tolerance of 1*e −* 4 and verified that the product of the map matrix with the null-space is several magnitudes smaller than the tolerance (< 1e *−* 11).

I computed the transformation matrices for force, equilibrium, and stability condition separately for each trial. The transformation map for equilibrium and stability does not rely on the muscle activations or any subject-specific measurements. By using nominal or cadaveric measurements to compute the maps, I am also able to ascertain how well the standard measurements reflect the real world behavior.

#### b. Alignment of the subspace obtained from task mechanics with the principal components

To find the alignments between the task relevant and irrelevant subspaces and the principal components, I first sorted the null-space and the complement space of the transformation matrices, T_efs_, T_e_, T_f_, and T_s_ by the decreasing amount of variance explained. A principal component analysis was performed on the muscle activations of each trial. I predicted that the first null-space component is aligned with the first principal component, where the task-irrelevant directions are ordered in decreasing values of fraction of the explained total variance. The cosine of angle, cos ϕ, between the first principal component and the task irrelevant direction that contains the maximal variance is calculated. The probability distribution of the angle between two random vectors in the seven-dimensional space using Monte Carlo simulation with 10^6^ samplings (figureS2) is used to report the p-values.

#### c. Analysis of change in null space with change in posture

For the task of producing the fingertip forces using the index finger, I calculated how the task irrelevant space changes with changes in posture. With a varying metacarpophalangeal (MCP) angle *θ*_1_ *∈* [−45°, 90°], proximal interphalangeal (PIP) angle *θ*_2_ *∈* [0°, 90°], and distal interphalangeal (DIP) angle *θ*_3_ *∈* [0°, 90°], I analyzed 9072 postures and calculated the intersection of the null spaces of T_f_, T_*e*_, and T_*s*_, to obtain the task irrelevant subspaces. To calculate the force map independent of the muscle electromyograms, I used the equation, T_f_ = J^−T^RF_iso_. The first null space vector of the task irrelevant subspace corresponding to the posture 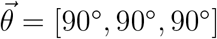 is selected as the reference direction to measure the change in angle of the null space vectors for other postures. Mean and the standard deviation of the angles are reported.

### E. Simplifications and assumptions

For the analysis in this study, I ignore the configuration dependent forces on the index finger. The rationale is that at 2N of distal fingertip force, the joint torques due to the constraint are [0.1264, 0.0384, 0.0130] Nm, whereas the joint torques due to gravity are [−5.5, −7, −1] × 10^−3^ Nm at MCP, PIP, and DIP joints, respectively. Thus, the joint torques due to gravity are two orders of magnitude smaller than that due to the constraint at the lowest force of 2N considered during the experiments. Ignoring the gravity terms is valid because a planar finger in contact has only one degree of freedom. For a 1D system, the null-space is continuous and would vary only by a small amount if gravity is considered [22]. Therefore, I ignore the terms due to gravity in further analysis and consider only the forces due to the constraint. For the muscle model, I ignore the force-velocity properties, the stiffness due to the force-length properties, and also ignore the short-range damping of the muscles.

### F. Parameter values and software

The matrix RF_iso_ is calculated from the muscle parameters obtained from cadavers [21, 23, 24], and phalanx lengths and masses used were measured during an earlier study [18, 25] (table S1). MATLAB R2022a Update 6 (9.12.0.2170939) was used for analysis.

## III. Results

### a. Force subspace

I confirm the previous findings [14] that the total variability is significantly lower in the task-relevant subspace than in the task-irrelevant subspace of the muscle activation to force mapping. The ratio of the task-relevant to task-irrelevant variability for the force mapping, *η*_f_, over 35 trials is 0.41 ± 0.05 (mean ± SE, figure 2) and significantly differs from 1 (t-test, *p* < 4*e* − 13, CI = (0.31, 0.52), figure 2).

**FIG. 1.**
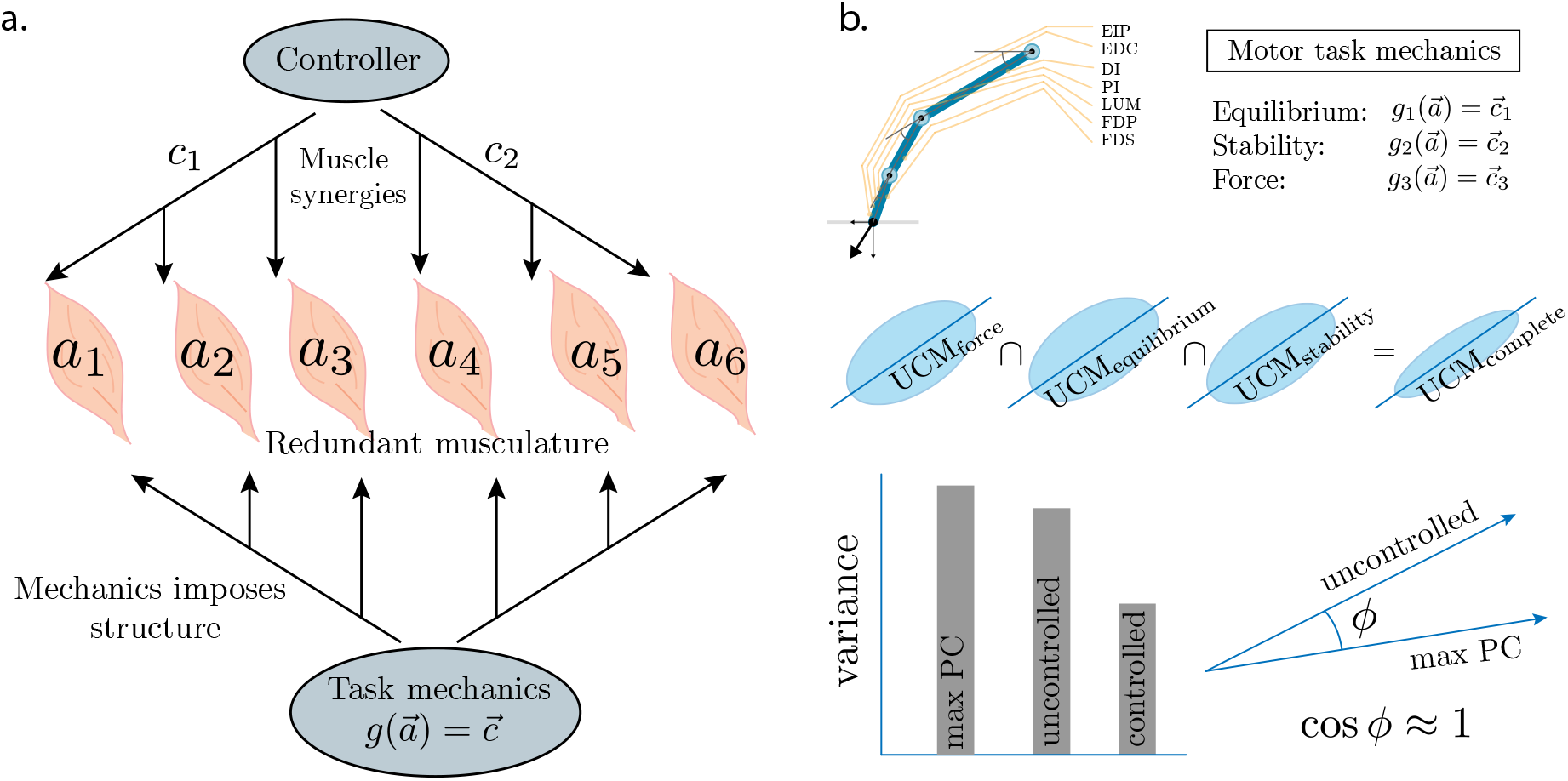
Pictorial representation of the hypothesis and the analysis. **a**, Muscle synergies simplify the neuromuscular control of the redundant musculature for a motor task. The mechanical requirements of a motor task also impose a structure on the muscle activities of a musculoskeletal system. The total redundancy of the musculoskeletal system depends on the degrees of freedom available after satisfying the mechanical requirements. **b**, For the success of a static motor task of producing instructed fingertip forces while maintaining a posture, muscle activities must satisfy the mechanical requirements of equilibrium, stability, and instructed force. Such demands are investigated by constructing individual uncontrolled manifolds for each mechanical demand and taking the intersection of the subspaces to derive an uncontrolled manifold that accounts for the complete mechanics of the task at hand.

**FIG. 2.**
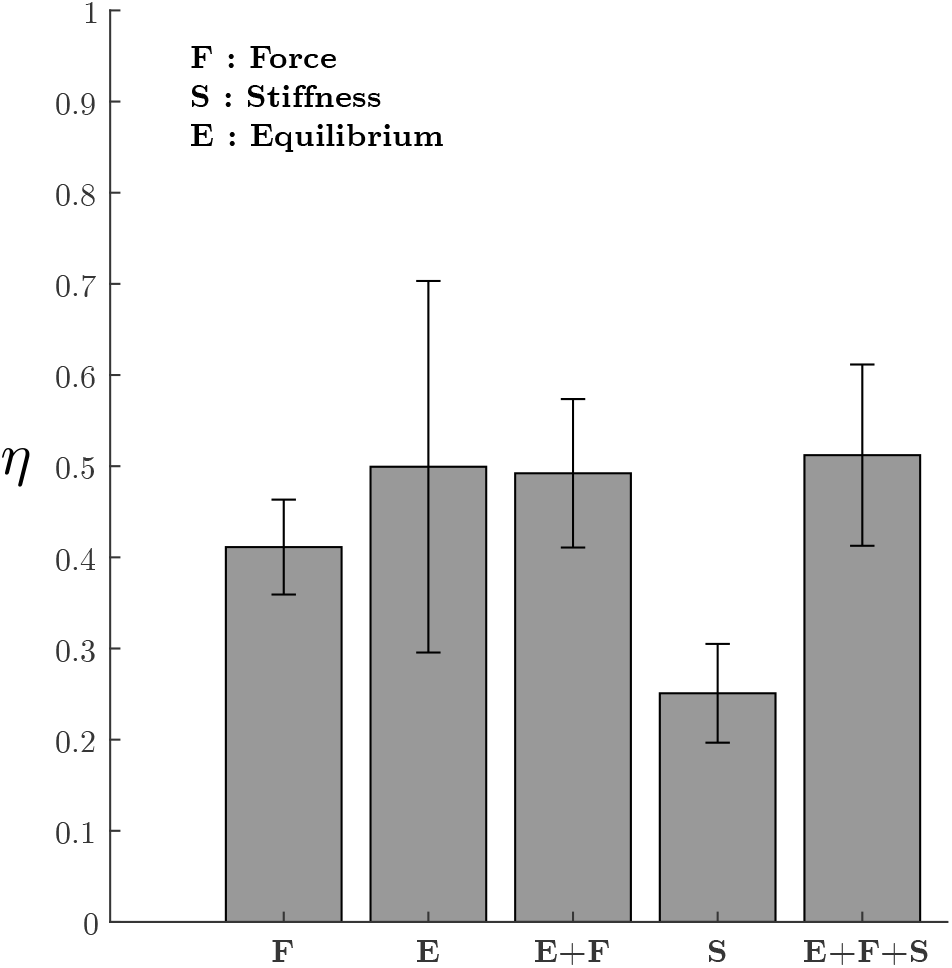
Descriptive statistics of the variability ratios for equilibrium, force, stiffness, and combined conditions. The height of the bars denote the mean, and the whiskers are the standard error of the mean.

### b. Equilibrium subspace

During the experiment, volunteers maintain postural equilibrium. The mean ratio of the task-relevant to irrelevant variability, *η*_e_, over 35 trials is 0.50± 0.20 (mean ± SE, figure 2), and it is significantly different from 1 (t-test, p = 0.02, CI =(0.09, 0.91), figure2). The ratio of variability channeled in the task-relevant subspace to that in the task-irrelevant subspace when both equilibrium and force are considered, *η*_ef_, is 0.49 ± 0.20 (mean ± SE), and is significantly different from 1 (t-test, p = 0.02, CI = (0.33, 0.66)).

### c. Stiffness subspace

The constraint of stability implies that variability in muscle activations must be channeled in the null space of muscle activations to the postural stiffness mapping. A finger in contact has only one degree of freedom specified by one eigendirection in terms of the joint angles. Muscle activations must stiffen this eigendirection for stability. Using the map T_s_ between muscle activations and the relevant stiffness eigendirection, I observe that the variability due to muscle activations is higher in the 6D null-space of T_s_ compared to its 1D orthogonal complement. The ratio of task-relevant to irrelevant variability, *η*_s_, is 0.25 ± 0.05 (mean ± SE, figure 2) and is significantly different from 1 (t-test, p < 1e − 15, CI = (0.14, 0.36), figure2).

### d. Combined subspace

I find that the variance in muscle activations is preferentially channeled in the null space of the mapping between muscle activities to the complete task-relevant subspace. For all subjects, I obtain a 2D task-irrelevant subspace and a 5D task-relevant subspace. The mean ratio of the task-relevant to task-irrelevant variability *η*_efs_ is 0.51±0.10 (figure 2), and is significantly different from 1 (t-test, *p* < 2*e−*5, CI = (0.31, 0.71), figure2).

### e. Alignment of subspace obtained from task mechanics with the principal components

Task-relevant and irrelevant directions obtained from task mechanics are linked to low and high variances in muscle excitations, respectively. The principal component analysis is an alternative method of projecting the large dimensional information in muscle excitations to a low dimensional subspace explaining large variance in the data. I find that the principal components explaining the largest variance aligns closely with the task-irrelevant directions obtained from task mechanics (figure 3). The alignment between the first principal component and the first component of the null space of T_efs_, T_e_, T_f_, and T_s_ is 47.0° ± 2.8° (p=0.07), 40.6° ± 2.2° (p=0.03), 40.1° ± 2.4° (p=0.03), and 41.3° ± 1.9° (p=0.03), respectively. In comparison, the alignment of the first principal component with the task-relevant direction containing the lowest variance is almost orthogonal (figure 3), as expected for two random vectors in a high-dimensional space.

**FIG. 3.**
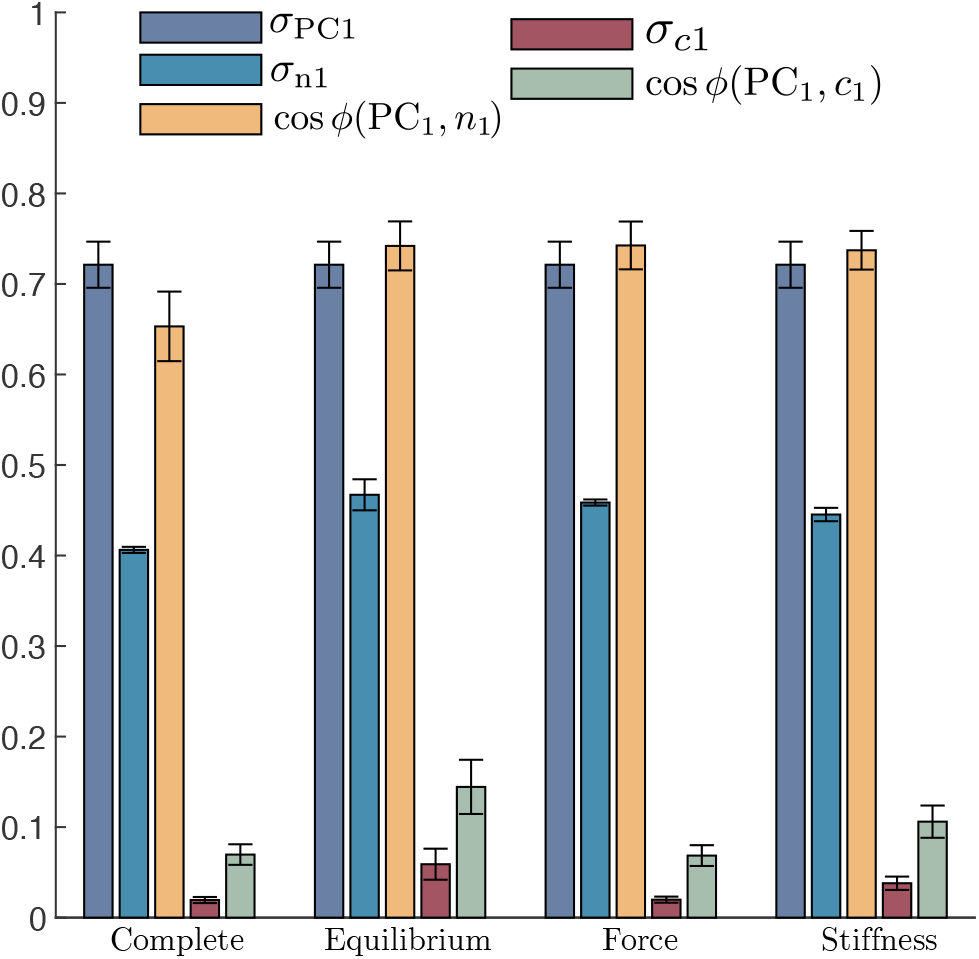
Cosine of the angle (cos(*ϕ*)) between the first principal component (PC1) and the task-irrelevant space vector 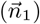 containing the highest projected fraction of total variance, and with the task-relevant space vector 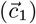 containing the lowest projected fraction of total variance, for the complete description of mechanics, and equilibrium, force, and stiffness conditions. *σ* denotes the fraction of total variance contained in either the first principal component, or the maximal relevant or irrelevant directions, as denoted by the subscript. The height of the bars represent the mean and the whiskers denote the standard error of the mean.

### f. Changes in the task irrelevant space with changes in posture

When varying the index finger posture through its approximate range of motion (SII), the first null space component of the complete task space containing the force, equilibrium, and stability conditions changes by 59.0° ± 20.5° (mean ± standard deviation).

## IV. Discussion

I show that the muscle activities support the minimum intervention principle of motor control when the task is to apply the instructed fingertip forces while maintaining postural equilibrium and stability. The relevant to irrelevant variability is consistently less than one for equilibrium, force, and stability conditions alone and when considered together to describe the complete task mechanics. People need not solely rely on muscle short-range muscle stiffness for postural stability at sub-maximal forces because joint damping may slow down the timescale of buckling instability, and sensorimotor feedback control may aid stability [18]. During the experiments analyzed in this study, the distal fingertip forces are sub-maximal, varying between 5 − 10 N, compared to 30 − 80 N in [18]. Yet, the results suggest that people likely rely on short-range muscle stiffness even at sub-maximal forces by activating muscles in a manner that preferentially control the muscle short range stiffness in the pertinent eigendirection prescribed by the minimal stiffness condition. Only select combinations of muscle activation patterns can help the finger stay in equilibrium, maintain stability, and produce the desired fingertip forces. I find five task-relevant directions and a two-dimensional null space in the muscle activation space of the complete map for the motor task of producing fingertip forces. Thus, an index finger in contact has a degree of redundancy of two after meeting the mechanical requirements.

The null space vector with the highest projected variance aligns with the principal component that explains maximal variance. Dimensionality reduction of muscle activities during a variety of motor tasks show that a few principal directions explain most of the variance, which are interpreted as task-relevant directions and invoked as evidence of muscle synergies [19, 20, 26–28]. The results of the current study suggest that the first few principal directions that explain most variance likely reflect the task-irrelevant directions, whereas the principal directions that explain low variance describe the task-relevant directions or the dimensions of the control space (figure 3). Thus, principal directions with large variances have consequences for understanding the level of muscle redundancy. Furthermore, the task-irrelevant null space changes by 59° when the phalanx angles span their range of motion, suggesting that the redundant directions and the principal component explaining the most significant variance must show a corresponding change when the posture changes. Whether muscle synergies for the control of redundant musculature only exist in the task-irrelevant subspace, and if muscle synergies facilitate or are organized independently of mechanics are open questions.

Modeling studies in biomechanics and motor control assume that muscle activations satisfy the mechanical constraints of equilibrium and stability [14, 18, 21]. Maintaining equilibrium and stability is necessary for performing motor tasks successfully; however, the biomechanical models used for describing motor tasks are approximate and rely on several simplifications and assumptions. For example, limbs are modeled as links with ideal joints, ligament stiffness, muscle contraction dynamics, and tendons properties are ignored, and constant instead of frequency-dependent muscle short-range stiffness and damping are assumed. Furthermore, the parameter values used for muscle moment arms, physiological cross-sectional area, optimal lengths and tension, phalanx masses and dimensions, and scaling constant of short-range muscle stiffness with tension mostly rely on the measurements from human cadavers or other animals and are not subject-specific. Despite these limitations, biomechanical models generate experimentally verifiable predictions suggesting that modeling assumptions do not cause information loss [18, 29]. Here, I provide direct evidence that muscle activities satisfy the modeling assumptions of equilibrium and stability, and also show the validity of using cadaveric measurements.

## V. Acknowledgements

Madhusudhan Venkadesan (MV) and Francisco J Valero Cuevas (FVC) for sharing data. MV and Ali Yawar for discussions and comments on the earlier manuscript drafts. FVC for pointing out some of the relevant literature. Department of Mechanical Engineering and Materials Science, Yale University, for providing shelter during my Ph.D. when I was conducting the above research.

