## Supplemental File 1 for "The minimum intervention principle of optimal control relates the uncontrolled manifold to muscle synergies": ucm_supplement.pdf

Neelima Sharma*[1,2]

[1] *Department of Organismal Biology and Anatomy, University of Chicago, Chicago, IL, USA*
[2] *Department of Mechanical Engineering & Materials Science, Yale University, New Haven, CT, USA*

### Contents

**S1 Derivation of the map from muscle activations to stiffness** **2**

**S2 Random noise does not generate the features of uncontrolled manifold** **3**

**References** **6**

### List of Tables

Table S1 Muscles used for actuation and their properties . . . . . . . . . . . . . . . . . . 5

### List of Figures

Fig. S1 Random noise does not generate the features of uncontrolled manifold . . . . . 3
Fig. S2 Distribution of the angle $\phi$ between two random vectors in the positive quadrant of a 7D space. . . . . . . . . . . . . . . . . . . . . . . . . . . . . . . . . . . . . . . 4

*

### S1 Derivation of the map from muscle activations to stiffness

The minimal stiffness matrix required for the stability of the finger posture is given by [1],

$$K_m = -\frac{\partial J^T \vec{f}}{\partial \vec{\theta}}\bigg|_{\vec{\theta}_0, \vec{\tau}_0} \tag{S1.1a}$$

$$\vec{f} = (JM^{-1}J^T)^{-1}JM^{-1}\vec{R}F_{iso}\vec{a} = GRF_{iso}\vec{a} \tag{S1.1b}$$

$$\text{where } G = (JM^{-1}J^T)^{-1}JM^{-1} \tag{S1.1c}$$

I project the dynamics on the null space to the constraint using the projection matrix $P = \vec{v}\vec{v}^T$ where $\vec{v} = \text{null}(J_0)$ and expand the constraint force $\vec{f}$ in terms of muscle activations,

$$PKP = -P\frac{\partial J^T GRF_{iso}\vec{a}}{\partial \vec{\theta}}P \tag{S1.2}$$

I rewrite the expression for the minimal stiffness matrix in the Einstein notation to obtain

$$P_{ij}K_{ji}P_{ij} = -P_{ij}\frac{\partial J^T_{jk}G_{km}R_{ml}F_{iso,ll}\vec{a}_l}{\partial \vec{\theta}_i}P_{ij} \tag{S1.3a}$$

$$K'_{ij} = -Q_{imj}R_{ml}F_{iso,ll}\vec{a}_l \tag{S1.3b}$$

$$\text{where } Q_{imj} = P_{ij}\frac{\partial J^T_{jk}G_{km}}{\partial \vec{\theta}_i}P_{ij} \tag{S1.3c}$$

Because the system is one-dimensional, the matrix $K'_{ij}$ has only one finite eigenvalue $\zeta_m$, and the corresponding eigenvector is $\vec{v}$. Similarly, the matrices $Q_{i1j}, Q_{i2j}, Q_{i3j}$ have only one finite eigenvalue given by $\zeta_{q1}, \zeta_{q2}, \zeta_{q3}$, with the corresponding eigenvector $\vec{v}$. I rewrite the equation (S1.3b) in terms of the relevant stiffness direction,

$$\vec{k}_m = \zeta_m\vec{v} = [\zeta_{q1}\vec{v}, \zeta_{q2}\vec{v}, \zeta_{q3}\vec{v}]RF_{iso}\vec{a} \tag{S1.4a}$$

$$\implies \vec{k}_m = T_s\vec{a} \tag{S1.4b}$$

$$\text{where } T_s = [\zeta_{q1}\vec{v}, \zeta_{q2}\vec{v}, \zeta_{q3}\vec{v}]\,RF_{iso} \tag{S1.4c}$$

### S2 Random noise does not generate the features of uncontrolled manifold

I provide a proof based on the Monte Carlo approach to the statement that fitting a linear model in the presence of random noise in the activation signal does not automatically generate an uncontrolled manifold. I constructed a mean force vector $\vec{f} = (1, 1, 10)$ N and added a uniformly random noise $\in [-1, 1]$ to generate a vector of length $10,000$ to simulate a time varying noisy signal. Relying on the observation that people do not drastically modulate their muscle activities during the task, I define the mean activation vector $\vec{a} = (0.3, 0.4, 0.5, 0.9, 0.9, 0.3, 0.4)$ and added a uniform random noise $\in [-0.1, 0.1]$ to generate a vector of length $10,000$ to simulate a time-varying noisy signal. Using the time varying vectors, I find T such that $\vec{f} = \mathrm{T}\vec{a}$. I find the complement space and null space of the mapping T, projected the variance in constructed muscle activities on the complement and the null space, and used it to calculate the variability ratio as done in Methods. For $10000$ Monte Carlo simulations, I show a histogram of the variability ratio and find that it is centered around 1. I separate the instances when the variability ratio is less than one and when it is greater than or equal to one. Observing figure S1, I conclude that random noise cannot generate an uncontrolled manifold.

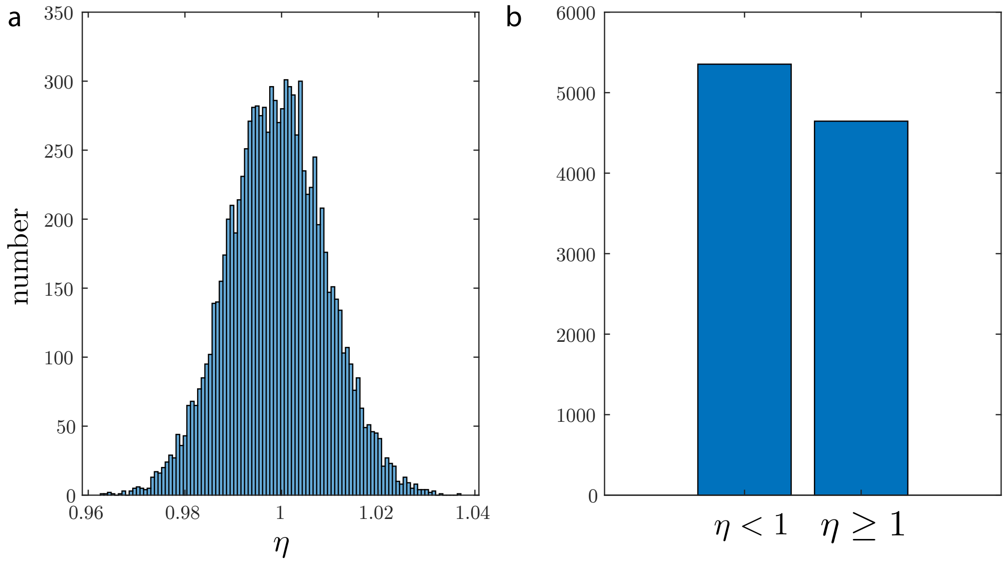


**Fig. S1:** **a.** Histogram of the variability ratio. **b.** Variability ratio separated by instances when it is less than 1, and when it is greater than or equal to 1.

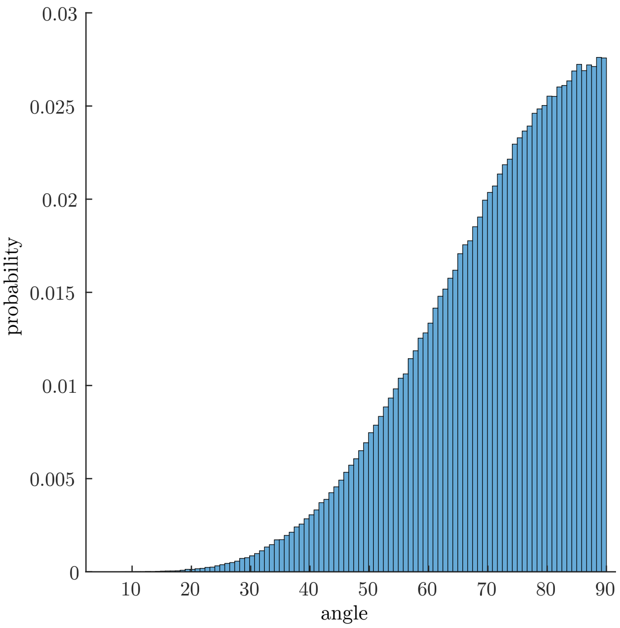


**Fig. S2:** Distribution of $\phi$ between two random vectors in the positive quadrant of a 7D space. Two vectors in a 7D space have a high probability of being orthogonal to each other. I use this distribution to compute the p-values of obtaining a given angle between two vectors.

**Table S1: Muscles used for actuation and their properties [1].** Seven musculotendon units, *flexor digitorum profundus* (FDP), *flexor digitorum superficialis* (FPS), *extensor indicis proprius* (EIP), *extensor digitorum communis* (EDC), *first lumbrical* (LUM), *first dorsal interosseous* (DI), and *first palmar interosseous* (PI) - actuate the three joints of the index finger, namely metacarpophalangeal joint (MCP), proximal interphalangeal joint PIP and distal interphalangeal joint (DIP) actuate the index finger joints.

| | FDP | FDS | EIP | EDC | LUM | DI | PI |
|---|---|---|---|---|---|---|---|
| Moment arms at finger joints, R (mm) [2, 3] | | | | | | | |
| MCP joint | 12.00 | 13.20 | -7.77 | -7.77 | 7.00 | 2.00 | 4.00 |
| PIP joint | 6.50 | 5.85 | -2.75 | -2.75 | -2.75 | 0 | -2.75 |
| DIP joint | 3.64 | 0 | -1.50 | -1.50 | -1.50 | 0 | -1.50 |
| Muscle physiological properties | | | | | | | |
| PCSA ($cm^2$) [2, 3] | 4.10 | 3.65 | 1.12 | 1.39 | 0.36 | 4.16 | 1.60 |
| Muscle tension, $t_i$ (N) [2, 3] | 143.50 | 127.75 | 39.20 | 48.65 | 12.60 | 145.60 | 56.00 |
| Optimal lengths, $l_i$ (cm) | 7.5 [4] | 8.4 [4] | 5.9 [4] | 7.0 [4] | 4.9 [5] | 2.9 [6] | 2.9 [7] |
